# Modification and validation of a GAD-GFP mouse line without accelerated aging-related hearing loss

**DOI:** 10.1101/2025.03.29.646110

**Authors:** Kaley Graves, William Dai, Kaitlyn Ortgiesen, Daniel A. Llano

## Abstract

GABAergic neurons in the inferior colliculus (IC) play a crucial role in auditory processing by extracting specific features of sounds (Ono et al., 2005). The Gad67-GFP mouse model developed by Tamamaki et al. in 2003 on a Swiss background facilitates studying these neurons by using a green fluorescent protein that is expressed endogenously via the GAD67 promoter. Unfortunately, this mouse suffers from accelerated aging-related hearing loss, limiting its utility in studying the auditory system. Here, we report the results of an 8-generation backcross of this line onto CBA/CaJ mice, which produces mice with stable low-threshold hearing while retaining GFP expression in GAD+ neurons. Additionally, this study investigates mechanisms that underlie hearing loss in the Gad67-GFP mouse model by focusing specifically on cochlear hair cells (HCs) and ribbon synapses, which may contribute to both model-specific hearing loss and clinical disorders like presbycusis. Findings revealed the newly generated F1 mouse model that resulted from the Gad67-GFP x CBA/CaJ backcross maintained better hearing thresholds when compared to ABR data for Gad67 and Swiss mice and very closely resembled those of the CBA/CaJ mice, mirroring progression of presbycusis in humans. Additionally, all morphological changes observed in cochlear structure correlated to ABR thresholds. F1 mice continued maintained expression of the GAD67 promoter in the IC via immunostaining.

## INTRODUCTION

Hearing loss affects 15% of the total adult population in the United States, which translates to roughly 38 million people over the age of 18 years old (Hoffman et al., 2017). Of those 38 million people, presbycusis, or age-related hearing loss (AHL), is the most common cause of hearing loss and is most often seen in an elderly population. Previous studies have shown that the risk of presbycusis doubles in each decade of life before eventually affecting as much as 50% of elderly adults by the age of 70 or older (Lin et al., 2011a). As aging occurs, integration of multiple sound stimuli in daily listening environments slows due to lapses in peripheral and central auditory processing, which can lead to cognitive deficits and loss of speech intelligibility.

The current study is designed to modify a mouse model that not only functionally examines GABAergic inhibition in the inferior colliculus (IC) but also allows mice to retain largely normal hearing throughout the lifespan. A GAD-GFP mouse model developed in 2003 by Tamamaki et al. has been instrumental in helping to understand the physiology of GABAergic neurons throughout the brain (Tamamaki et al. 2003). Unfortunately, it is available on a Swiss background strain that demonstrates accelerated aging-related hearing loss. This study also explores potential cause of hearing loss in this animal, with a strategy to correct its hearing loss by backcrossing it onto a CBA/CaJ strain, thus permitting its broader use to study GABAergic neurons in the auditory system.

Mouse models are commonly used for hearing research for many reasons, such as the fact that the mouse genome is 80% like the human genome and has been fully sequenced, which makes it much easier to determine a more specific genetic deficit that could be an underlying cause of hearing loss (Pennacchio, 2003). Three of the four mouse models featured in this study have been identified as having rapid progressive sensorineural hearing loss (SNHL) or age-related hearing loss (AHL or presbycusis).

The first model, C57BL/6J (C57), typically exhibits elevated hearing thresholds that are consistent with severe hearing loss at roughly 9-12 months of age (Kikkawa et al., 2012). *CDH23* has been identified as a primary mutative gene that causes progressive SNHL in the C57 mouse and has been genetically mapped as associated with hearing impairment in laboratory mice (Noben-Trauth et al., 2003). The second model chosen, the Swiss-Webster mouse, is known to have age-related SNHL that results from progressive degeneration of hair cells in the cochlea as well as a loss of spiral ganglion neurons (SGNs) and has a mutation in the *CDH23* gene (Henry & Chole, 1980; Shnerson et al., 1981, Willot et al., 1998; Drayton et al., 2006). The third mouse line, the Gad67-GFP knock-in mouse, is used to study cortical projections in the auditory cortex to and from the IC as a mechanism of studying top down and bottom up processing. While the Gad67-GFP knock-in mouse allows for visualization of auditory projections from the IC, it has elevated hearing thresholds early in its lifespan because of its Swiss-Webster background. The last control mouse line used in this study, CBA/CaJ, is most frequently used in hearing research because SNHL hearing loss is slow to progress and is more applicable to the human lifespan when assessing AHL (Frisina et al., 2011). Additionally, CBA mice do not have the known *CDH23* genetic mutation that C57 mice have that causes such a rapid progression of SNHL.

In the Drayton et al. (2006) study, the authors indicate that mouse strains that contain the *CDH23* gene without mutation are more likely to maintain life-long hearing, which includes the CBA/CaJ mouse strain that is of interest in this study. Strains that have the mutative *CDH23* gene, such as C57 and BALB/cJ, are more likely to experience early onset hearing loss that is rapidly progressive (Drayton et al., 2006).

Ultimately, the goal of this study is to create a mouse model that retains GFP expression in the IC under control of the endogenous Gad67 promoter while improving hearing thresholds, as measured by ABR. Morphological analysis will assess cochlear hair cell loss in all four strains of mice at 12 months of age. Results obtained from the current study are expected to better inform how cochlear structure affects hearing loss. Since mice share similar genomic mapping as humans and undergo controlled gene expression, this study aims to translate murine findings to human AHL. With presbycusis affecting roughly 50% of adults by age 75, identifying structural deficits may help clarify its progression in the cochlea and in the IC.

## MATERIALS & METHODS

### Animals

All experiments were conducted with adult glutamic acid decarboxylase-67 (Gad67)-GFP knock-in mice initially developed by Tamamaki et al. (Tamamaki et al. 2003) that are bred with CBA mice to obtain eight generations of mice of both sexes unless noted otherwise. Additional Swiss Webster mice and CBA/CaJ mice were purchased from The Jackson Laboratory (Bar Harbor, ME, USA). Founder Gad67-GFP mice were generously provided by Drs. Doug Oliver and Deborah Bishop from the University of Connecticut and are not commercially available for purchase. For this study, the first generation (F1) were used as the primary basis of comparison. All procedures were approved by the Institutional Animal Care and Use Committee at the University of Illinois. Animals are housed in care facilities that are approved by the American Association for Assessment and Accreditation of Laboratory Animal Care. The number of animals used were kept to an absolute minimum to reduce any potential for suffering at all stages of this study.

### Auditory Brainstem Response testing

Animals were anesthetized at 6 months of age and again at 12 months of age with intraperitoneal acepromazine (2-3mg/kg), ketamine hydrochloride (100mg/kg), and xylazine (6mg/kg) before insertion of three subdermal electrodes. Electrodes were placed at the vertex of the skull (main electrode), one behind the right ear (reference electrode), and one behind the left ear (ground electrode). Sound stimuli was presented using an ES1 freefield speaker (Tucker-Davis Technologies) in an anechoic sound chamber, with waveforms generated by SigGen software. The output of the speaker was calibrated at the appropriate frequencies used for testing: white noise, 4kHz, 8kHz, 16kHz, 32kHz, and 40kHz. A microphone (Model 377A06; PCB) and a preamplifier (SV 12L; Svantek) was used to calibrate per frequency. Each stimulus was presented for 5ms total time, with 3ms of constant-amplitude sound and 1ms cosine-shaped ramps, at a rate of 21 Hz. Raw potentials were obtained with an RA4LI headstage, RA16PA preamp, and an RA16 Medusa Base station (Tucker-Davis Technologies), filtered between 100-5000 Hz and averaged at 512 traces/sound stimulus. Hearing thresholds were determined via Wave I generation in decreasing intensity as measured in decibels (dB). Observation of the peak of wave I at 80 dB served as a starting threshold measurement. If there was not a measurable wave I at 80 dB, the animal was determined as having no hearing at that frequency. A supplemental dose of intraperitoneal ketamine in saline (50mg/kg) was administered if mice began to respond to toe pinch during ABR testing. Upon completion of ABR testing, mice were removed from the sound chamber and returned to home cages to recover from anesthesia.

### Histology

After 12-month ABR data was compiled, mice were euthanized for temporal bone and brain collection for histology to analyze morphological changes in the HCs and ribbon synapses in addition to confirming expression of GFP in the brain. Mice were given a lethal intraperitoneal injection of ketamine (200mg/kg) and xylazine (6mg/kg) for euthanasia via transcardial perfusion. Mice were also euthanized via carbon dioxide asphyxiation followed by cervical dislocation to collect temporal bones. Both the temporal bones and the brain were postfixed separately in 4% paraformaldehyde (PFA) overnight at 4C. Temporal bones were decalcified in 0.5-mM ethylenediaminetetraacetic acid (Fisher Scientific, Waltham, MA, USA) for 48 hours. Samples were then stored in 10-mM phosphate buffered saline (PBS) (Sigma-Aldrich, St. Louis, MO, USA) at 4C until use. The cochlea was dissected into three turns of equal length and placed free-floating in PBS into a 48-well plate for immunostaining for HCs and ribbon synapses. The primary antibody incubation step was conducted at 37°C in a walk-in warm room on a rotating plate overnight, followed by three washes in PBS at 5 minutes each at room temperature (RT). Secondary antibodies were applied to each well containing a cochlear turn, and the plate was incubated for 2-4 hours in a dark box to avoid photobleaching of the samples on a rotating plate to ensure optimal distribution of antibodies. After incubation of secondary antibodies, samples were washed with PBS for 5 minutes each at RT, followed by application of Hoechst to label cellular nuclei in a dark box for 15-20 minutes at RT on a rotating plate. A final three washes of PBS for 5 minutes each at RT was conducted prior to mounting in an anti-fade Prolong Gold mounting solution to provide stability of the tissue and prevent photo-bleaching and coverslipping for imaging, as described in Montgomery and Cox, 2016; Cai et al., 2018. Antibody labeling of Myo7a was used to label all HCs (both OHCs and IHCs), Sox2 was used to label supporting cells (SCs), Ctbp2 was used to label pre-synaptic ribbons, GluR2 was used to label post-synaptic ribbons, and Hoechst was used to label all cellular nuclei to help assess morphological changes in the structures of the cochlea, as stated in RQ2. All antibodies used for cochlear immunostaining are given in Table 1.

**Table 1:** Antibodies used for cochlear immunostaining.

| Cochlear antibodies | Dilution | Product number | Company |
| --- | --- | --- | --- |
| **Mouse IgG1 anti-Ctbp2** | 1:500 | Cat#BDB612044 | BD Biosciences,<br>San Jose, CA |
| **Mouse IgG2a anti-GluR2** | 1:200 | Cat#MAB397 | Millipore,<br>Billerica, MA |
| **Rabbit anti-myo7a** | 1:200 | Cat#25-6790 | Proteus BioSciences<br>Ramona, CA |
| **Goat anti-sox2** | 1:100 | Cat#AF2018 | R&D Systems<br>Minneapolis, MN |
| **Alexa 488 goat anti-mouse IgG2a** | 1:1000 | Cat#A21131 | Invitrogen,<br>Waltham, MA |
| **Alexa 568 goat anti-rabbit IgG** | 1:1000 | Cat#A11036 | Invitrogen,<br>Waltham, MA |
| **Alexa 647 goat anti-mouse IgG1** | 1:1000 | Cat#A21240 | Invitrogen,<br>Waltham, MA |
| **Hoechst trihydrochloride** | 1:2000 | Cat#H3570 | Invitrogen,<br>Waltham, MA |

To verify GFP expression in the IC, brain tissue was cryoprotected in an ascending series of sucrose solutions to preserve the integrity of the tissue prior to being embedded and cut into 40μm thick slices on a freezing sledge cryostat. Immunostaining for Gad67 in the IC was conducted as previously described (Lesicko et al., 2016). Sections were immunostained for Gad67 at 1:1000 using a mouse anti-Gad67 monoclonal antibody (cat#MAB5406, Millipore, Billerica, MA, USA). Secondary antibody immunostaining was conducted at 1:100 with Alexa 568-conjugated goat-anti mouse (cat#A11004, Invitrogen, Waltham, MA, USA), followed by three washes in PBS. Sections were mounted and coverslipped with an anti-fade solution to prevent deterioration of antibody labeling (Vectashield, Vector Laboratories, Newark, CA, USA). Brain histology was conducted to answer RQ3.

### Imaging and Analysis

To better assess what changes, if any, are observed in the morphology of the cochlea, imaging of the cochlear structures (the HCs and synaptic ribbons) was conducted using a Zeiss LSM 900 Airyscan confocal microscope. HCs were quantified using Zen Blue Lite (Zeiss) software. Morphological analysis of the OHCs were imaged at 20x to obtain a broad region of all three rows of OHCs, whereas IHCs containing the ribbon synapse pairs were imaged at 63x using the Airyscan module to obtain z-stacks for better resolution and visualization of matched pre- and post-synaptic ribbons through the different layers of tissue in cochlear whole mounts. Tile scans of sections of the IC were obtained at 5x to visualize the entire structure. Further adjustments for color balance will be done using Zen Blue Lite and Adobe Photoshop was used to draw masks around the edge of the sections to cover irrelevant brain tissues that were not needed for analysis of expression of GFP in the IC.

### Statistical Analysis

ABR data was analyzed via two-way ANOVA in OriginPro to determine significance across frequency and strain for threshold. Threshold outliers were identified using standard deviation and significance level of 0.05 via the OriginPro Outlier ID plug in for each frequency per strain. Tukey’s post-hoc test was used to determine comparisons between means for significance. Quantification of HCs were conducted via two representative 200μm regions per cochlear turn, per mouse strain. A one-way ANOVA was conducted with Excel. Ribbon synapses were quantified via two sections of 4-6 representative IHCs per turn per strain (N=3-4) and analyzed using a one-way ANOVA.

## RESULTS

### ABR Measurements

ABR Wave I peaks were used to determine thresholds of mice across each of the four groups because they are much more easily visualized in mice compared to wave V, which is used in humans (Zheng et al., 1999) (**Fig. 1-3**). White noise was used to determine that each mouse had functional hearing prior to beginning the ABR, since it is non-frequency specific sound stimuli. Additionally, wave I peaks represent the response from the acoustic nerve as stimuli travels from the peripheral inputs in the inner ear to the structures of the central auditory pathway.

**Fig. 1.**
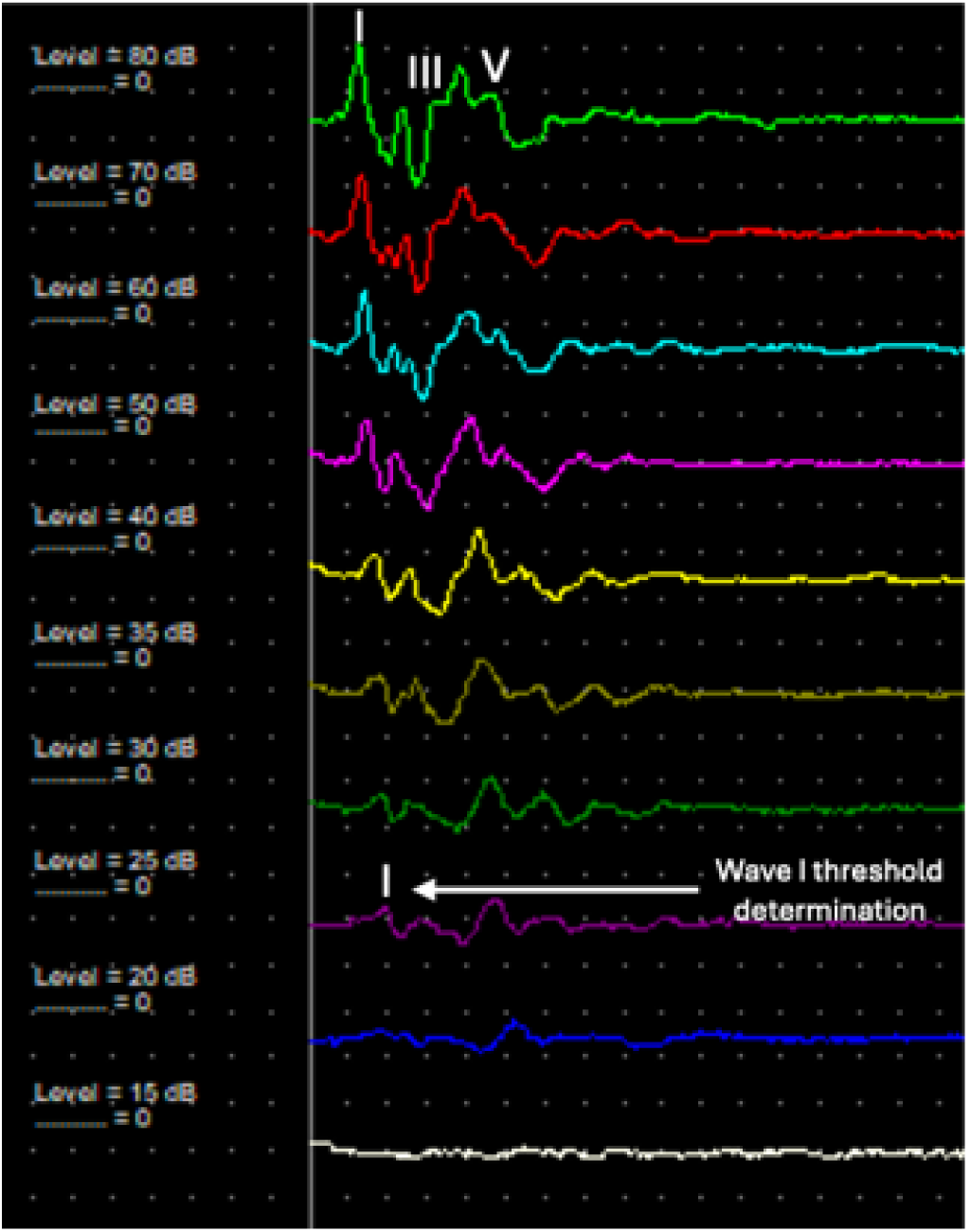
Example of waves I, III, V as shown in mouse ABR data in white noise and how ABR threshold is determined by presence of wave I in descending intensity.

**Fig. 2.**
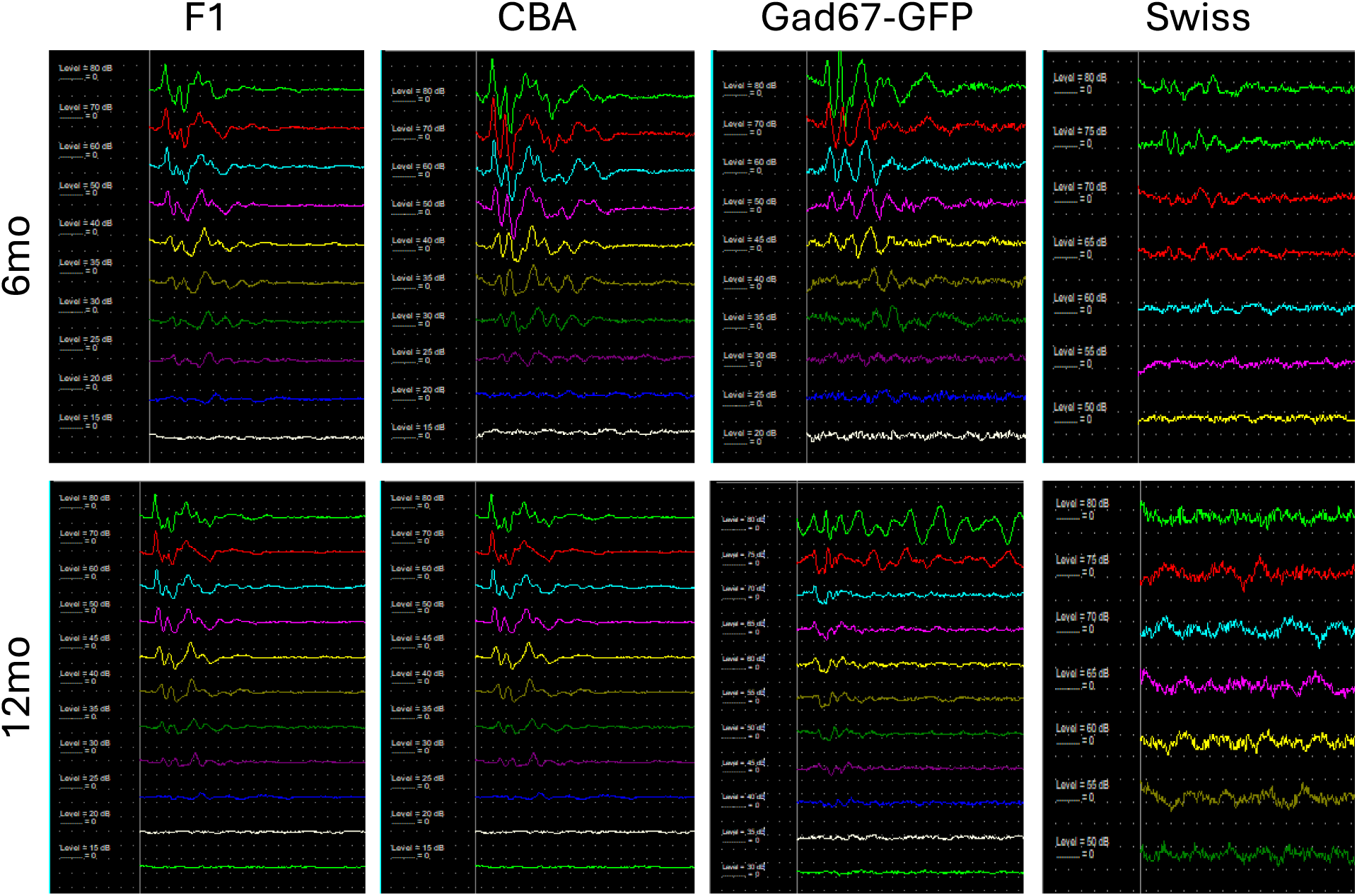
ABR waveforms at 6mo and 12mo across F1, CBA, Gad67-GFP, and Swiss mice in white noise to determine presence of functional hearing.

**Fig. 3.**
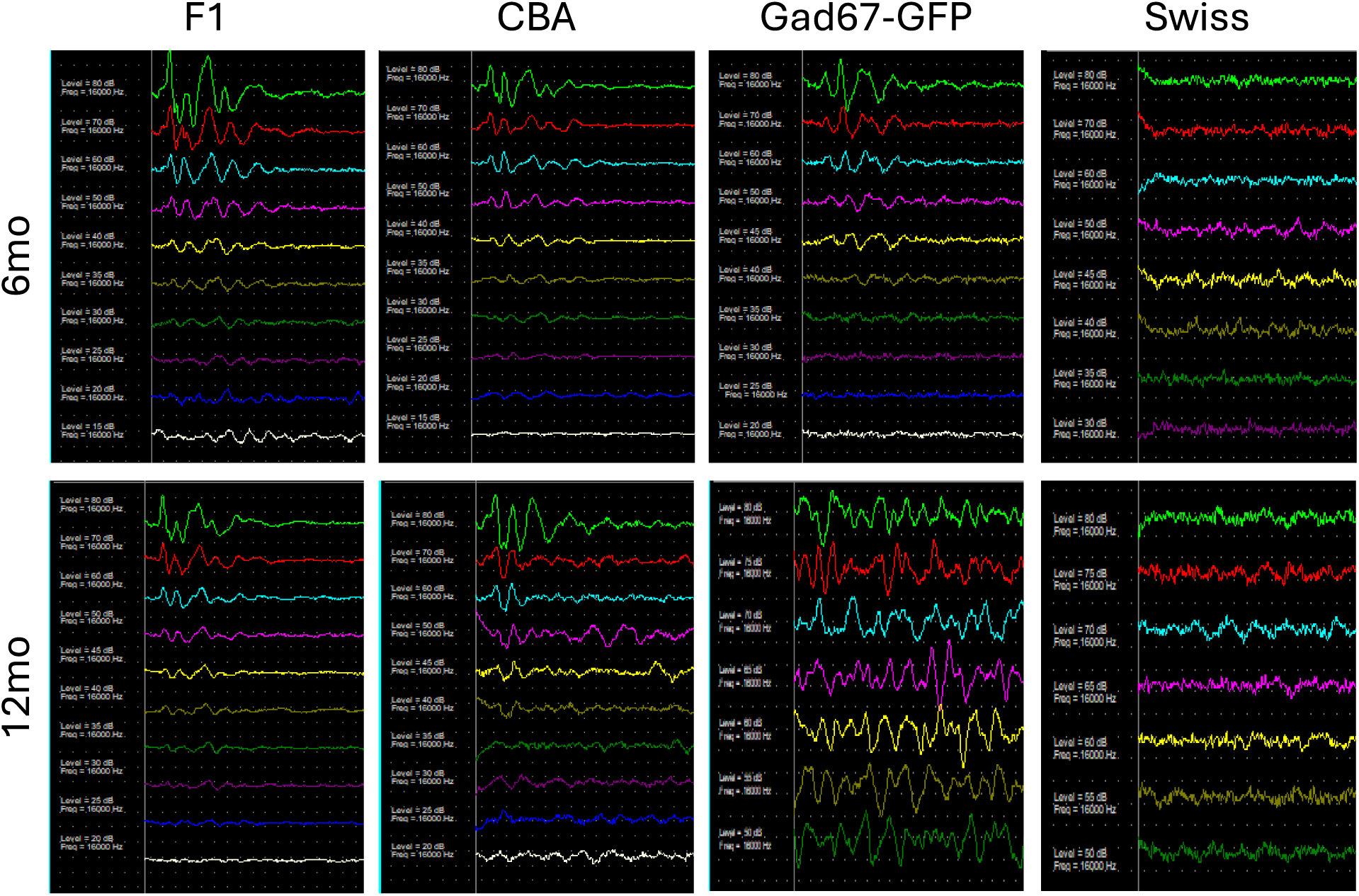
ABR waveforms at 6mo and 12mo across F1, CBA, Gad67-GFP, and Swiss mice at 16kHz.

A comparison of the ABR wave I threshold determination of the F1 generation of mice to the three control groups (CBA/CaJ, Gad67-GFP, Swiss) at 6 months of age shows that the F1 mice have consistently improved ABR hearing thresholds across all six frequencies (**Fig. 4**). A two-way ANOVA indicated that the interaction between both strain and frequency was significant (p<0.0001). However, only the interaction between 32 and 40 kHz was not significant when analyzed via Tukey’s post hoc, likely due to similar ABR thresholds.

**Fig. 4.**
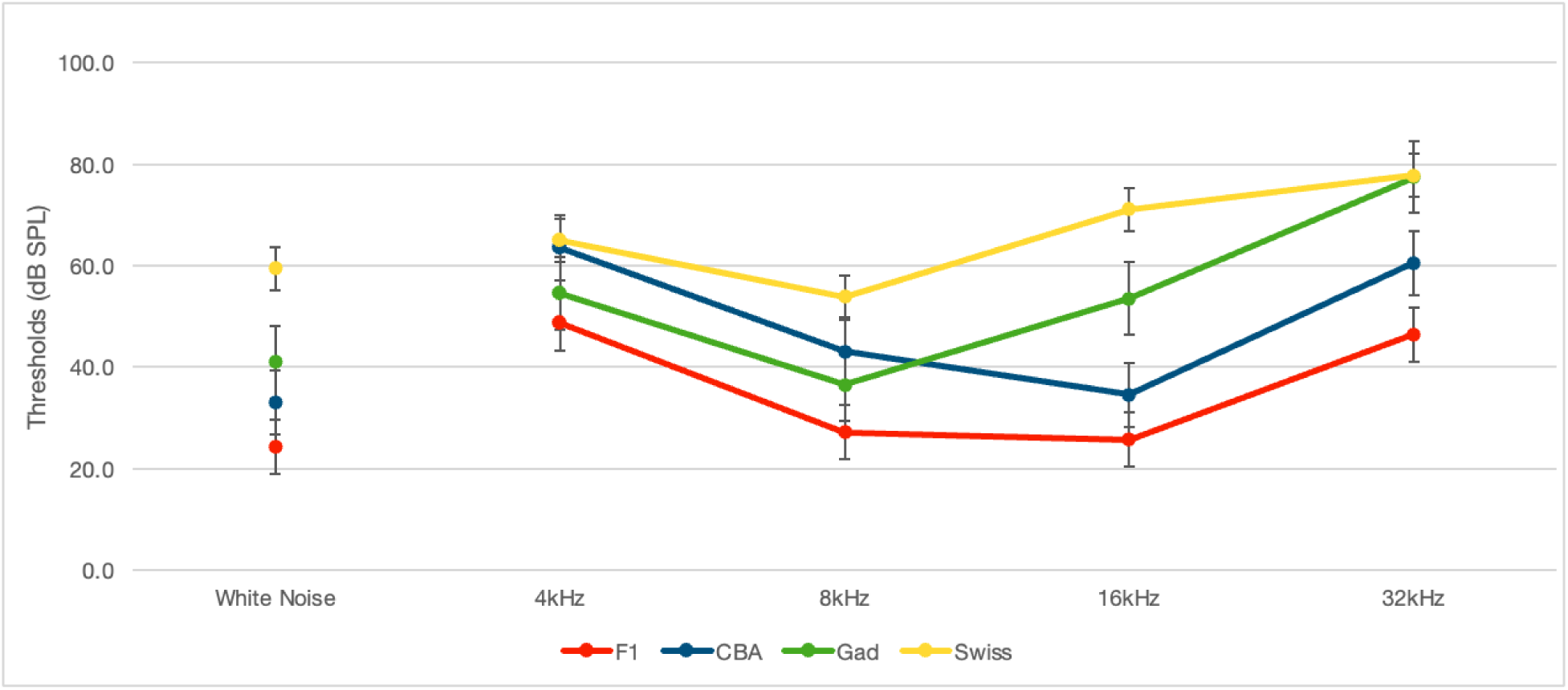
Average ABR thresholds at 6 months of age in F1, CBA, Gad67-GFP, and Swiss mice.

When ABRs were conducted again at a 12-month timepoint, F1 mice largely retained their hearing from 6 months of age and only exhibited slightly elevated thresholds. The most notable degradation of hearing occurred in the Gad67-GFP and Swiss Webster mice, which were almost completely deaf by the 12-month age measure (**Fig. 5**). A two-way ANOVA revealed that ABR thresholds were all statistically significant (p<0.001) when compared per frequency and strain, which supports age-related increases in thresholds at 12-month timepoints. Tukey’s post hoc indicated that there were significant interactions between all strains except Gad67 and Swiss, which supports Gad67-GFP being bred on a Swiss Webster background and exhibiting poor hearing at an early age. When analyzed for frequency, Tukey’s post hoc indicated significance between 4-8, 4-16, 8-32, 16-32, 8-40, and 16-40 kHz. This finding correlates with observed increased ABR thresholds at 12 months of age.

**Fig. 5.**
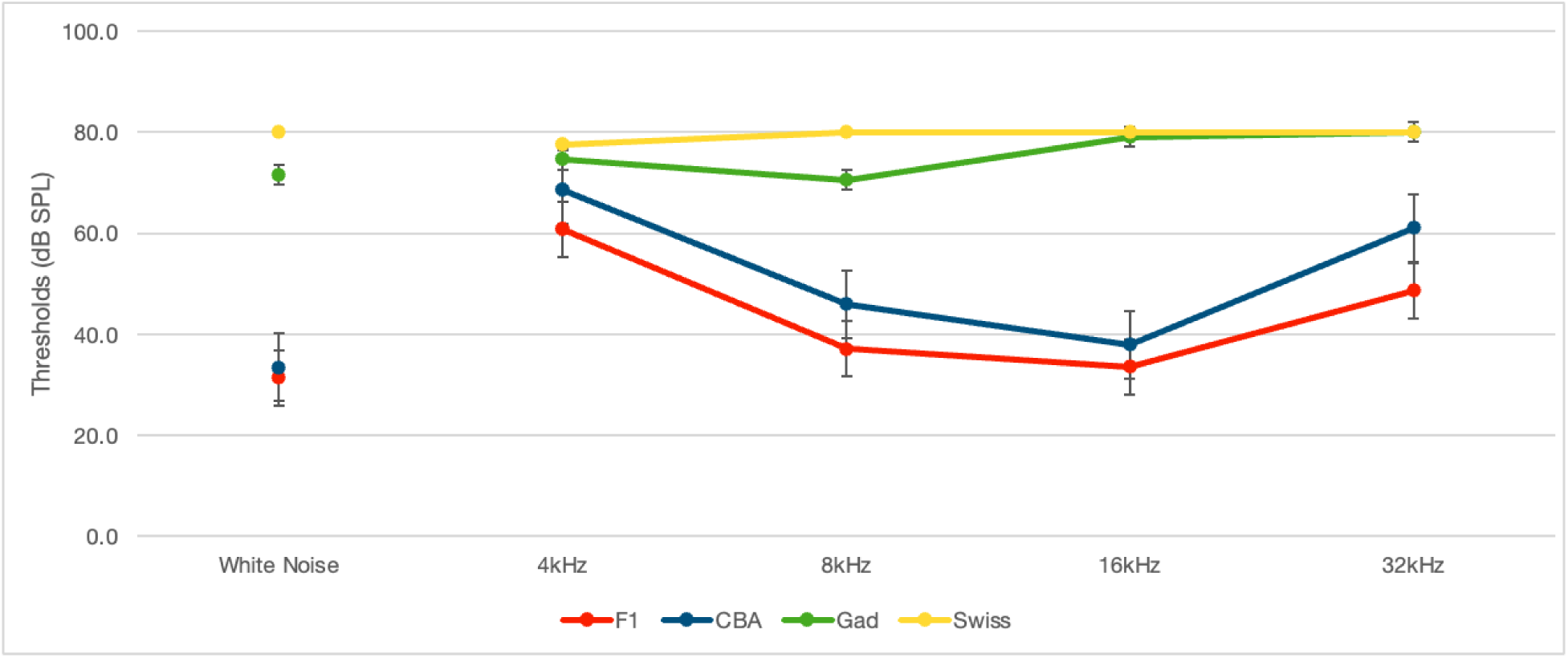
Average ABR thresholds at 12 months of age in F1, CBA, Gad67-GFP, and Swiss mice.

### Brain histology for GFP

Another important goal of the backcross is to retain endogenous GFP expression in the IC for ongoing studies in the Llano lab that look at cortical projections in auditory processing. Immunostaining for Gad67-GFP in slice physiology of F1 mice compared to Gad67-GFP control mice shows that eGFP expression is maintained. It should be noted that the Gad67 antibody does not label cell bodies very well and instead labels Gad67 positive neuropil, whereas endogenous GFP expression labels all cell bodies and not necessarily neuropil. The GAD modules contain both cell bodies and neuropil, which is why double labeling proves positive via immunostaining and endogenous expression (**Fig. 6**).

**Fig. 6.**
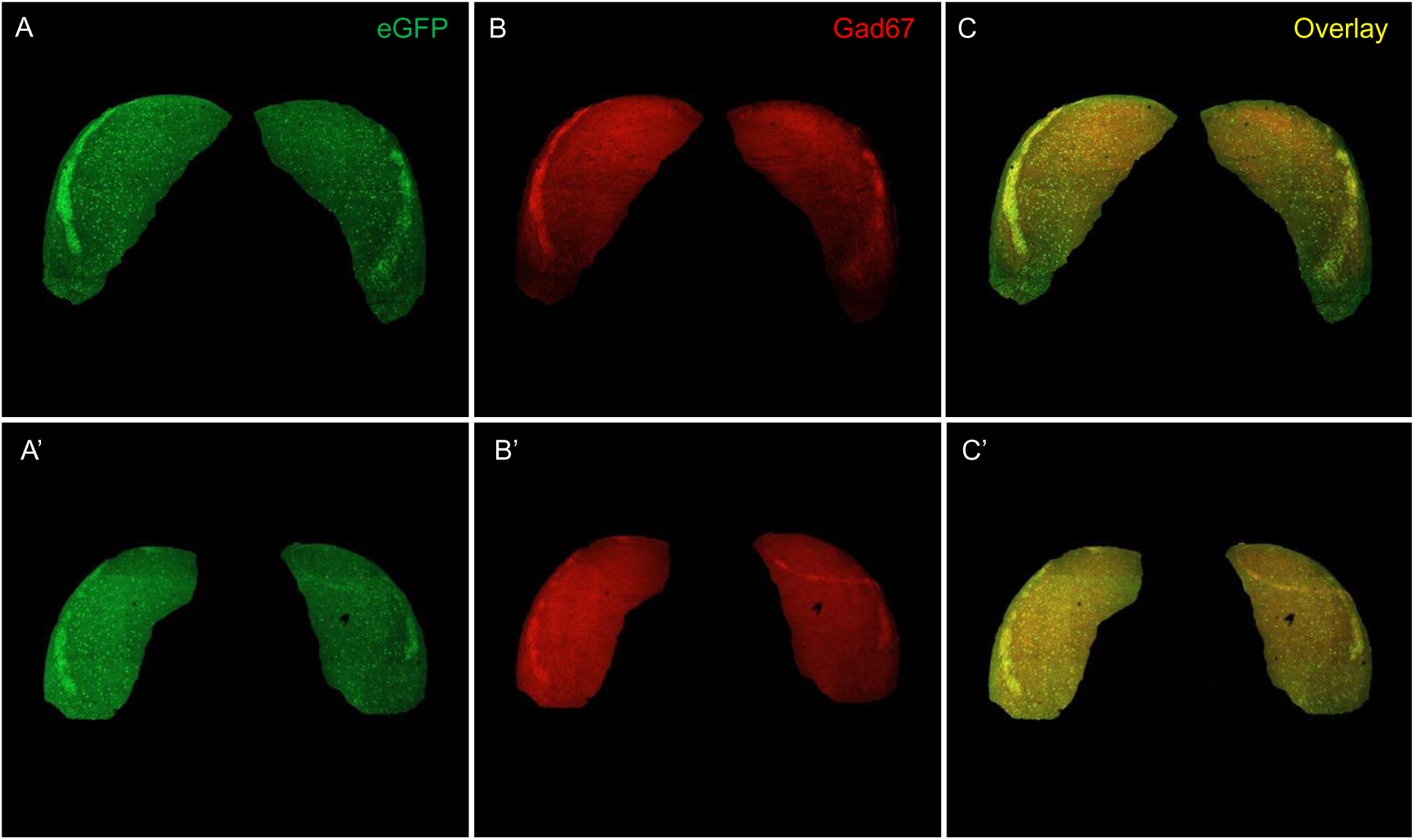
Endogenous GFP expression in the IC of the Gad67-GFP mouse (A-C) is maintained in the F1 familial generation backcross (A’-C’). Immunostaining for Gad67 (red) colabels with eGFP (green) to verify true Gad67 labeling. Debris was masked in each image shown to better demonstrate colabeling.

### Cochlear Morphology

The most crucial component of understanding possible correlated changes in hearing thresholds as measured via ABR lies in assessing cochlear morphology. Immunostaining for myo7a to label all OHCs and IHCs, sox2 to label supporting cells, and Hoechst for cellular nuclei across all four mouse strains revealed notable differences in cochlear structure. Sox2 was used as a secondary cellular label to differentiate between a HC and a SC to avoid potential miscounting. Quantification of OHCs and IHCs was conducted on two randomly chosen 200μm representative sections per each turn of the cochlea. When analyzing cochlear morphology to answer RQ2, there were notable differences in the structure of the Gad67-GFP and Swiss mice compared to the F1 generation and CBA mice. The three rows of OHCs are linear and well organized, with an occasional missing HC or an extra HC (**Fig. 7**) in the CBA and F1 mice but are almost missing entirely in the Gad67-GFP and Swiss mouse strains. This correlates with ABR threshold findings since Gad67-GFP and Swiss both demonstrate an almost complete loss of hearing by 12 months of age across most frequencies tested. Further confirmation of a lack of HCs in the Gad67-GFP and Swiss mice is seen in Sox2 staining (**Fig. 8**) and displays the disorganization of SCs while supporting the lack of any visible HCs. Quantification revealed minimal differences in the number of OHCs and IHCs between F1 and CBA mice in all three turns of the cochlea, with no quantification conducted on any of the Gad67-GFP and Swiss mice due to missing HCs. A one-way ANOVA indicated that quantification for OHCs, and IHCs, was statistically significant (p<0.001). This finding correlates with observed increased ABR thresholds at 12 months of age and supports presbycusis as a result of morphological changes in the cochlea.

**Fig. 7.**
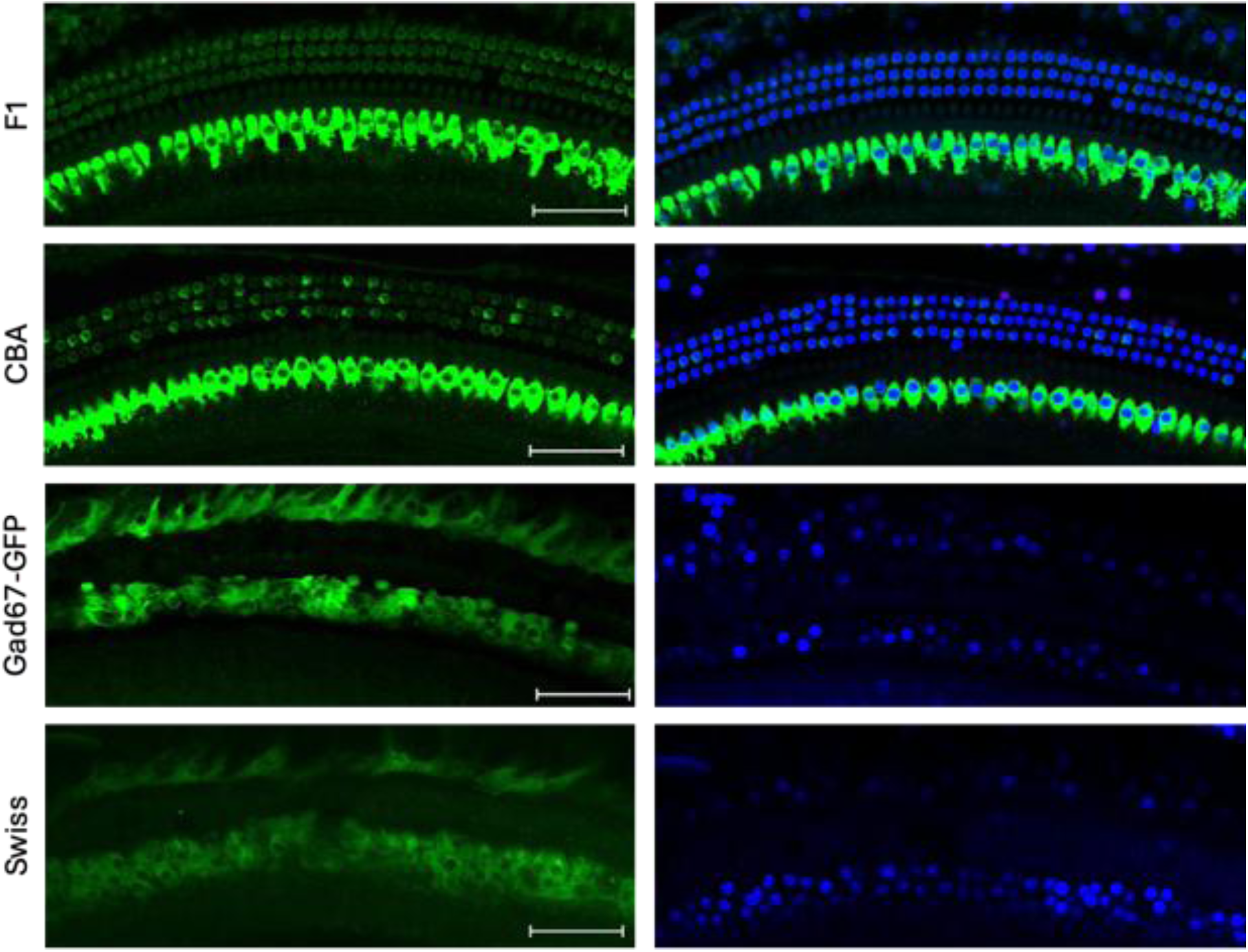
Observed morphological changes across all four strains in mice in the middle turn, as labeled with myo7a (green) and confirmed with a Hoechst nuclear stain (blue). N=3, scale bar = 50µm.

**Fig. 8.**
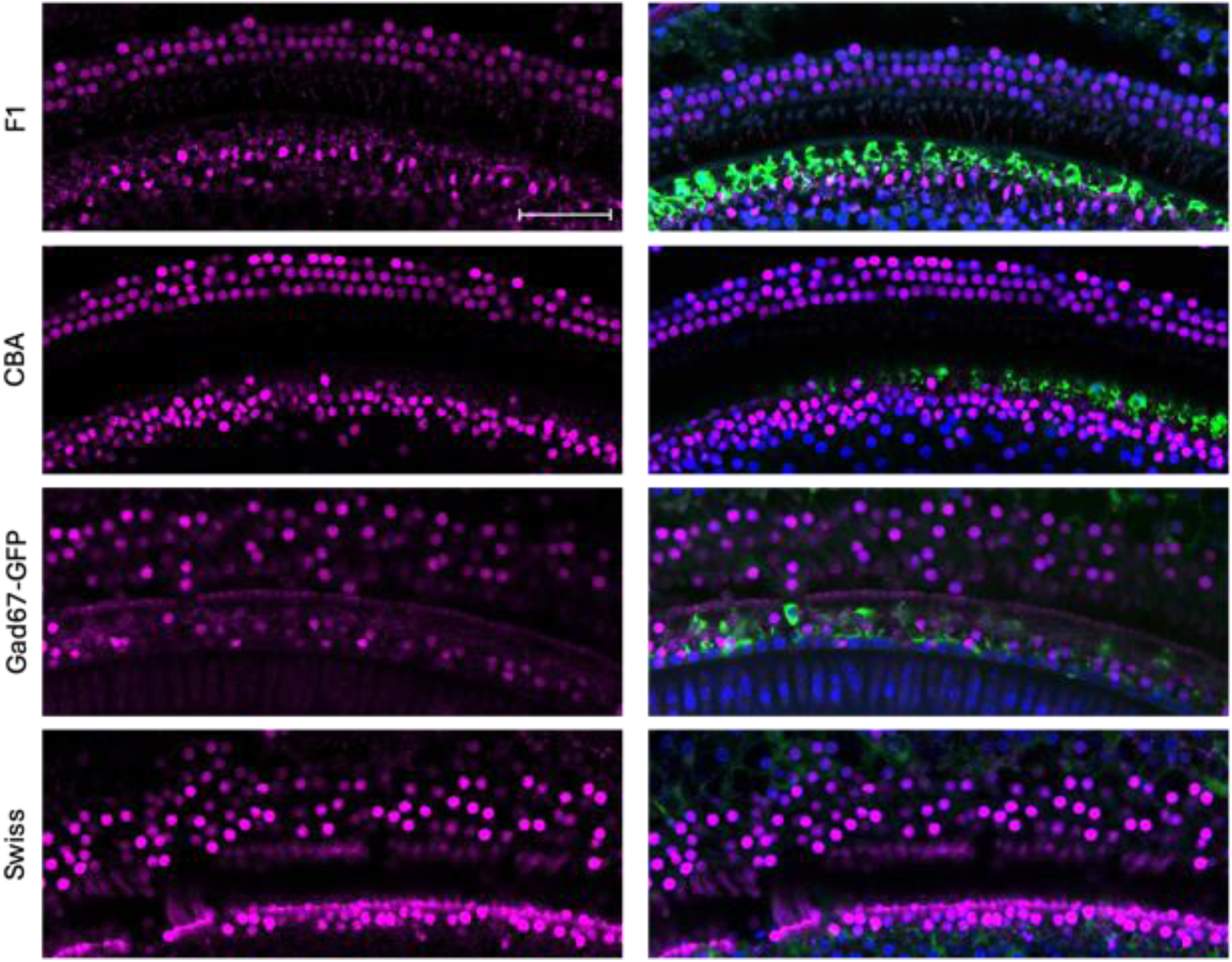
Observed morphological changes across all four strains in mice in the middle turn, as labeled with Sox2 (magenta) and confirmed with a Hoechst nuclear stain (blue). Some HC labeling observed in F1 and CBA mice with myo7a (green). N=3, scale bar = 50µm.

In addition to observed lack of HCs (both OHCs and IHCs) in Gad67-GFP and Swiss mice, IHC distribution was observed to be lacking in the basal turn, occasionally sparse but present in the middle turn, and somewhat present in the apical turn in the CBA and F1 mice. This finding correlates with determined ABR thresholds when considering frequency location in the cochlea. Quantification of ribbon synapses across two chosen representative sections comprised of 4-6 IHCs imaged at 63x were significant (p<0.05) across all four strains when compared for the total number of pre- and post-ribbon synapse pairs. (**Fig. 9**)

**Fig. 9.**
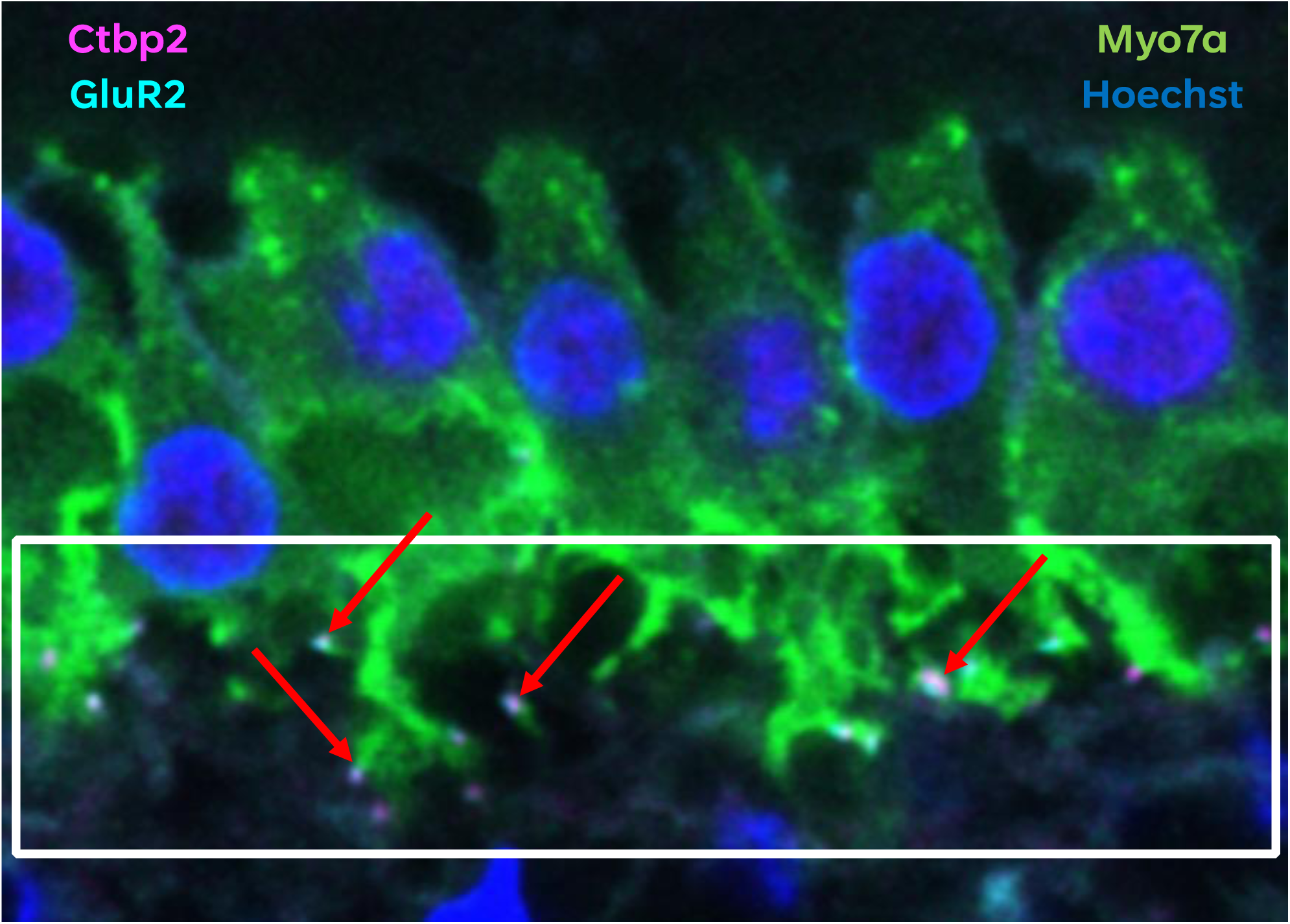
Observed pre- and post-ribbon synapse pairs in IHCs, as labeled with Ctbp2 (magenta), GluR2 (cyan), myo7a (green), and a Hoechst nuclear stain (blue). Matched pre- and post-ribbon synapses are indicated by red arrows and appear as white due to immunofluorescence overlap (N=3-4).

## DISCUSSION

### Generation of F1 mice exhibit better hearing thresholds

As shown previously, the F1 mouse strain maintains better hearing thresholds when compared to ABR data for Gad67-GFP and Swiss mice. These ABR thresholds are like those of the CBA mice, which exhibit steady progression of mild sensorineural hearing loss that closely resembles age-related hearing loss in humans. Elevated thresholds observed at 4 kHz at both timepoints across all four strains of mice are not easily explained despite validation and equipment calibration. However, this is consistent with other studies that investigated potential confounds to explain elevated 4 kHz thresholds during ABR analysis. Sha et al. (2008) did not observe any accompanying HC loss in the apical turn of the cochlea at 3 months of age, which is when elevated 4 kHz thresholds began to appear via ABR. It was suggested that further analysis of this cochlear pathology could be conducted via ribbon synapse quantification or determining changes in stereocilia bundles on the HCs, since there are no visible morphological changes to the HCs themselves (Sha et al., 2008).

Another previous study has shown that an F1 generation resulting from a CBA x C57 backcross has improved hearing thresholds, as determined via ABR (Frisina et al., 2011). This is due to only having one copy of the *ahl* gene, which requires one copy to be present for phenotypically normal hearing. In our study of F1 mice resulting from a Gad67-GFP x CBA backcross, our findings suggest that there is also only one copy of the *ahl* gene present due to the introduction of the CBA parent strain with the Gad67 (which has a shared C57 background with the Gad67-GFP and Swiss mice).

Additional secondary control F1 generation ABR testing conducted in both Gad67-GFP positive and Gad67-GFP negative mice revealed nearly identical hearing thresholds at 6 months of age, which is consistent with the original 6-month findings in only Gad67-GFP positive F1 mice. When compared to age-matched CBA mice at 6 months that were tested at the same time, all thresholds followed the trend seen in our previous findings. ABR testing is ongoing in this F1 generation of GFP positive/negative mice with age-matched CBA controls to confirm similar thresholds at a 12-month timepoint.

### Observed Differences in Cochlear Morphology

HCs were shown to be almost entirely absent in aged Gad67-GFP and Swiss mice but were still organized neatly with minimal missing cells or abnormalities in F1 and CBA mice. Other studies have shown that CBA/CaJ mice maintain hearing thresholds with a slow progression of high frequency hearing loss and minimal HC loss until roughly 12mo of age and that there are significant differences in the number of HCs present when compared to C57 mice (Sha et al., 2008). Park et al. (2010) also investigated other structures in the cochlea, such as the stria vascularis and SGNs, to account for morphological changes beyond the HCs that could account for age related hearing loss. The findings in our study corroborate previous research studies in that sensorineural presbycusis is likely a result of missing HCs or lack of HC function, but we cannot exclude the possible contribution of other morphological changes that may occur in the cochlea.

Interestingly, even though the number of OHCs far outweigh the total number of IHCs in the cochlea, the IHCs are primarily responsible for transmission of acoustic information to the brain via ribbon synapses. Quantification of pre- and post-ribbon synapse pairs across all four mouse strains further supported ABR threshold determination and were statistically significant when compared across groups. Post-mortem research in humans has shown that cochlear synaptopathy can often occur well before any loss of HCs is seen, contributing to presbycusis (Viana et al., 2015). In other species such as gerbils, ribbon synapses are lost primarily in the apex of the cochlea in old age before progressing to the middle and basal turns, compared to an observed loss in the apical turn of CBA mice (Gleich, Semmler, & Strutz, 2016). Conversely, this study observed more loss of ribbon synapse pairs in the basal turn, which is consistent with C57 mice, as evidenced by the quantification of ribbon synapses in the Gad67-GFP and Swiss mice that are bred on a C57 background. However, Gleich et al. determined that the loss of ribbon synapses was not necessarily related to presbycusis in gerbils and suggested that endocochlear potential (EP) is more relevant to maintain sensitivity of the inner ear to sound stimuli (2016). Unfortunately, EP measurement is challenging to obtain in mice due to the small size of the cochlea and limited space for microelectrode insertion through the tympanic membrane, as well as the limited volume of the perilymph (Ohlemiller, Hartsock, & Salt, 2022). As a result, other structures of the cochlea certainly cannot be overlooked for analysis in future studies assessing cochlear morphology. However, regardless of cochlear synaptopathy or HC loss observed in previous studies, our study demonstrates that morphological changes in cochlear structure correlate to ABR thresholds.

### Maintained expression of GFP in the IC

The Gad67-GFP mouse model is used in the Llano lab to trace GABAergic neurons in the IC, so maintained endogenous GFP expression is crucial in the backcross with CBA when generating familial offspring. Our study confirmed that eGFP is robust in all F1 mice that are Gad67-GFP+. Subsequent familial generations also maintain robust expression, as assessed via slice histology of the IC. Access to easy neuronal tracing helps provide more information as to how auditory information that is transmitted from the peripheral auditory pathway to the central auditory pathway interacts with the auditory cortex in mouse models. According to Lesicko et al., GABA is found in a modular distribution in the lateral cortex of the IC (2016). However, the function of modular GABA is yet to be determined and is currently under investigation in the Llano lab. Unfortunately, the Gad67 antibody does not label cell bodies very well and instead labels the neuropil, which is also found in Gad67 modules. Neuropil are areas where there are dense networks of interwoven nerve fibers, along with branches, synapses, and glial filaments. Immunostaining in the backcrossed F1 mice show a phenotypically normal distribution of GABA in these modules, which is evidenced via endogenous GFP expression in addition to the red immunostaining for GAD.

### Limitations & Future Directions

ABR testing in conjunction with cochlear morphology indicates that hearing loss occurs due to a loss of HCs and/or ribbon synapses, but there are many other mechanisms in the cochlea that could also play a role. Further investigation into the correlation between ABR thresholds and morphology is necessary to strengthen the argument that presbycusis occurs due to loss of HCs and/or ribbon synapses. Distortion product otoacoustic emissions (DPOAE) testing is another way of assessing HC function in the cochlea and is used frequently in humans as well as in murine studies for a basis of comparison to ABR thresholds. Equipment to conduct DPOAE testing in mice was not available at the time of ABR testing in this study but would be a good secondary measure to strengthen the correlation between morphological changes in the cochlea and ABR threshold determination. Previous studies that compared C57 and CBA mice have shown that not only do C57 mice exhibit a greater loss of HCs at an early age when compared to CBA, but also that DPOAE testing confirmed ABR findings as a functional measure (Park et al., 2010). Furthermore, Frisina et al. (2011) assessed OHC functionality via DPOAEs in F1 hybrids of C57 x CBA mice and found that DPOAEs were significantly better preserved in old age than either the C57 or CBA parent strains and measured similarly to DPOAEs obtained at a younger age.

When assessing cochlear morphology, this study is limited in that we cannot look at all structures in the cochlea to better ascertain possible causes of age-related hearing threshold shifts. Confocal microscopy is limited to four fluorescent channels and additionally, some of the antibodies that could be used for immunostaining are made in conflicting host serum(s). Additionally, assessment of stereocilia bundles was successful when immunostaining but available confocal microscopy equipment did not have a high enough magnification that would provide high resolution images to ascertain any clear deficits or structural issues in each bundle. This could be remedied in the future by either obtaining a 100x objective or sending slides of cochlear turns to colleagues who have access to a scanning electron microscope, which would provide high resolution images of stereocilia bundle morphology. Liu et al. (2022) demonstrated that stereocilia and OHCs were lost at 22 months of age in CBA mice, while the majority of IHCs were still present but had elongated individual stereocilia. Based on current morphological analysis in this study, we would anticipate seeing similar structural deficits in mice that have present HCs.

Loss of all OHCs and IHCs seen in the Gad67-GFP and Swiss mice also could be confirmed via TUNEL (terminal deoxynucleotidyl transferase) assay to determine presence of apoptotic cells. Park et al. (2010) confirmed severe apoptosis in the basal turn of C57 mice as early as 6 months of age when compared to CBA mice. Further investigation of other cochlear structures, like SGNs and the stria vascularis, would provide a more comprehensive argument for presbycusis occurring as a result of morphological changes. Viana et al. (2015) stated primary neural degradation could contribute to presbycusis. In future animal studies, the connection between SGNs and ribbon synapses should be explored via immunofluorescence and confocal microscopy when there are elevated ABR thresholds and present synaptic pairs in the IHCs to determine if SGNs are another morphological contributor to presbycusis.

Despite the caveats of animal research, mice are still commonly used in hearing loss research due to structural similarities between the mouse and human inner ear. Gad67-GFP mice enables the study of GABAergic neuron function in the IC but develops early onset hearing loss due to a mutation in the *CDH23* gene from the Swiss-Webster mouse background. The F1 backcross with a CBA/CaJ mouse model offers an improved auditory profile while allowing maintained expression of eGFP. Additional investigation of cochlear morphology supports a close relationship between loss of HCs and/or ribbon synapse pairs and ABR thresholds, which suggests that the F1 mouse model could be an ideal model for future research in the central auditory pathway.

## Summary & Conclusion

In summary, presbycusis is one of the most prevalent causes of hearing loss in the elderly and affects roughly 50% of the population by the age of 70. However, animal models are more commonly used in age related hearing loss research due to structural similarities between the mouse and human inner ear that help better explain some of the fundamental biological processes that occur in hearing loss. Our mouse model that is used for studies involving GABAergic neurons in the IC permits study of GABAergic function via endogenous GFP expression in the Gad67-GFP mouse. However, this mouse loses hearing at an early age due to a mutation in the *Cdh23* gene from the Swiss Webster mouse background. We have developed an F1 backcross with a CBA/CaJ mouse strain that enables maintained expression of eGFP while improving hearing thresholds as determined via ABR testing at 6 and 12 months of age. Additional investigation of cochlear morphology supports a close relationship between loss of HCs and/or ribbon synapse pairs and ABR thresholds. Our F1 mouse model (Gad67-GFP x CBA) could be an ideal model for future research in the central auditory pathway with improved hearing that most closely resembles presbycusis in humans.

## Acknowledgments

This work was supported by NIDCD R01DC013073 and The Kiwanis Neuroscience Research Foundation.

## Notes

### Competing Interest Statement

The authors have declared no competing interest.

